# Evolutionary double-bind treatment using radiotherapy and NK cell-based immunotherapy in prostate cancer

**DOI:** 10.1101/2024.03.11.584452

**Authors:** Kimberly A Luddy, Jeffrey West, Mark Robertson-Tessi, Bina Desai, Taylor M. Bursell, Sarah Barrett, Jacintha O’Sullivan, Laure Marignol, Robert A Gatenby, Joel S Brown, Alexander RA Anderson, Cliona O’Farrelly

## Abstract

Evolution-informed therapies exploit ecological and evolutionary consequences of drug resistance to inhibit the expansion of treatment-resistant populations and prolong time to progression. One strategy, termed an evolutionary double-bind, uses an initial therapy to elicit a specific adaptive response by the cancer cells, which is then selectively targeted by a follow-on therapy. Here we examine the combination of radiation therapy and immunotherapy as a quantifiable double-bind strategy. Radiotherapy (RT) induces lethal double-strand DNA breaks, but cancer cells can adapt by upregulating DNA damage response pathways. While this evolutionary strategy increases resistance to DNA damaging agents, it also results in enhanced expression of natural killer (NK) cell ligands potentially increasing vulnerability to an immune response.

Using a radiation-resistant human prostate carcinoma cell line (22Rv1), we demonstrate that RT-resistant cells upregulate NK cell ligands, including major histocompatibility complex class I chain-related protein A/B (MICA/B), and poliovirus receptors (PVR1, PVRL2) with a 2-fold increase in sensitivity to NK cell mediated killing.

We investigated this potential evolutionary double bind through *in vitro* studies and evolution-based mathematical models. Radiotherapy alone slowed overall growth but strongly selected for RT-resistant cells. NK cell therapy alone suppressed the RT-resistant population but with a surviving population of radiation-sensitive cells. These dynamics were framed mathematically, and model simulation predicted optimal tumour control would be achieved through initial RT rapidly followed by NK-based immunotherapy. Subsequent experiments confirmed the model prediction. We conclude that radiotherapy and NK cell-based immunotherapy produces an evolutionary double bind that can be exploited in heterogenous tumours to limit RT resistance.

## Introduction

Evolution-informed therapies are novel therapeutic strategies with two key objectives: 1) reduce overall tumour burden while also 2) preventing the outgrowth of a fully resistant phenotype (1, 2, 3). Therapy resistance represents a fundamental barrier to curative treatment strategies. Resistance occurs when a population of tumour cells contain pre-existing traits or acquire adaptive traits enabling their survival during therapy. Evolution-informed strategies often exploit the fitness cost of this resistance. That is, the investment of cancer cell resources to produce, maintain, and operate the molecular machinery of resistance increases fitness when treatment is administered but decrease fitness in the absence of treatment (4, 5). The latter typically results in a proliferative disadvantage compared to non-resistant cells in the absence of treatment (4, 6). Furthermore, the molecular machinery itself can produce an increased sensitivity to a second treatment or alter the tumour cell’s interactions with the immune system (7). These dynamics can be exploited by generating an “evolutionary double bind” in which the mechanism(s) of resistance to the first therapy produce specific vulnerabilities that can be targeted by a subsequent treatment (6, 8, 9).

The evolutionary double bind strategy for treating cancer was derived from “predator facilitation” seen with owls and snakes (8, 10). Dessert rodents adapt to predation from owls by hiding. This increases the hunting efficiency of dessert snakes that reside in the brush (10)(Figure 1A). A double bind in cancer occurs when the first therapy facilitates the efficacy of the second therapy. Importantly, because of these dynamics, the second therapy may be highly effective in context of a double bind but much less so when given as an initial treatment.

**Figure 1.**
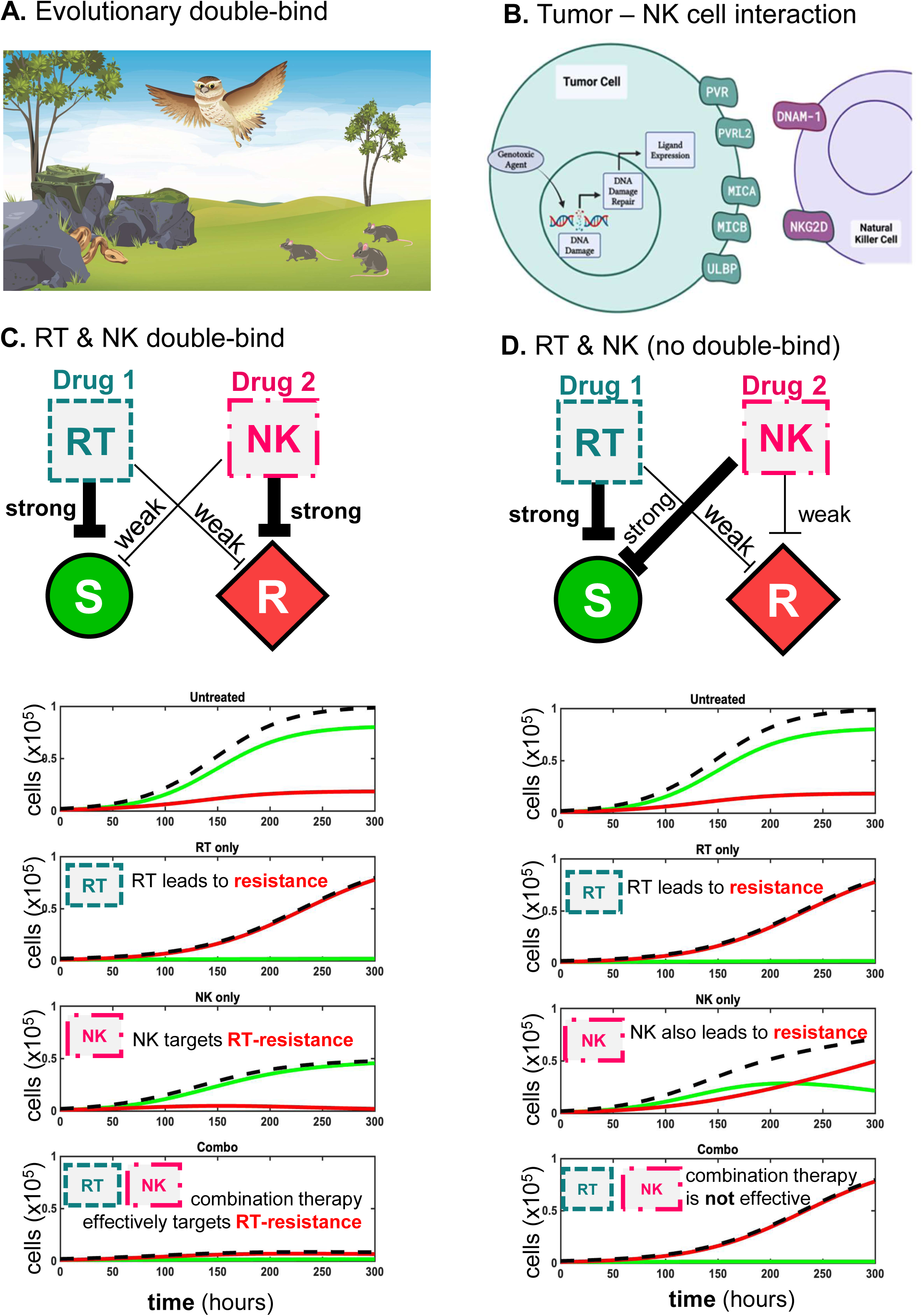
Evolutionary double bind therapy utilizes two therapies that explicitly target populations with different modes of therapy resistance. (**A**) The owl-snake dynamic is used to illustrate the ecological concept of an evolutionary double **(B)** NK cell ligands are expressed in response to genotoxic stressors including radiotherapy. **(C)** Hypothesized double-bind between radiation therapy (RT) and natural killer (NK) cells. RT strongly targets RT-sensitive cells (green) and weakly targets resistant cells (red). NK cells strongly target RT-resistant cells and weakly target RT-sensitive. Example treatment trajectories shown, where RT leads to RT-resistance, NK leads to RT-sensitivity, and combination effectively targets both cell types. **(D)** In contrast, without a double-bind, NK cells may strongly target RT-sensitive cells, rendering NK and combination therapy not effective in eliminating resistant cells.

Radiotherapy (RT) and immunotherapy are an appealing combination for cancer treatment (11). Given locally, RT induces double-stranded DNA breaks in tumour cells initiating cellular stress responses that trigger either cell death or DNA repair and cell survival (12). T cell-based immunotherapies benefit from RT-induced activation of the innate immune response and novel tumour antigens produced during RT that can generate a systemic immune reaction (11, 13). Radiotherapy also changes the protein expression on surviving cancer cells. Most notably, it has been shown that RT, as well as other DNA damaging agents, increases natural killer (NK) cell ligand expression on tumour cells (Figure 1B) (14, 15).

NK cells induce apoptosis in cancer cells with perforins and granzymes packed into cytolytic granules, and through the release of death-inducing ligands such as Fas ligand (FasL) and TNF-related apoptosis-inducing ligand (TRAIL) (16, 17). Additionally, activated NK cells support the adaptive immune response through the release of chemokines and cytokines including IFN-gamma, TNF, GM-CSF, IL6, CCL3, CCL4, and CCL5 (18). The activation of NK cells relies on a combination of inhibitory and activating signals received from target cells (19, 20). Activation of NK cells occurs only when these signals result in a net activation signal, that is, when activating proteins on the surface of tumour cells achieve a larger signal than regulatory proteins. Tumour cells often increase inhibitory signals (e.g., altering ligand expression on tumour cells) as an adaptive strategy for immune evasion (21, 22, 23).

Here, we hypothesize that the altered expression of NK ligands on tumour cells in a radiotherapy-resistant population would increase the sensitivity to NK cell mediated killing, creating an opportunity for an evolutionary double bind treatment strategy. Untreated conditions lead to a growth of both RT-sensitive (green) and RT-resistant (red) cells, where RT-sensitive cells outcompete RT-resistant cells due to a small cost of resistance. When radiation is applied, selection for RT-resistance occurs. In a true double-bind (left column, Figure 1C), NK cell-based therapy would select for radiation-sensitive cells. Seen in the schematic, this requires that NK therapy strongly inhibits radiation-resistant cells while weakly inhibiting sensitive cells. This criterion results in best response for combination therapy: inhibiting the growth of both sensitive and resistant cells (Figure 1C).

In contrast, the right column illustrates the scenario when the criterion for a double-bind is not met (Figure 1D). Here, NK cell-based therapy strongly inhibits radiation-sensitive cells and only weakly inhibits RT-resistant cells. Under combination therapy, radiation resistance is still under strong positive selection, illustrating the lack of an evolutionary double-bind without the strong-weak inhibition criterion.

Using an isogenic prostate cancer cell line model of radiation resistance derived from 22Rv1 cells, we show that NK receptor ligands are differentially expressed on the radiation-resistant cell line compared to the control line. We found that the radiation-resistant line was more sensitive to NK cell-mediated killing, indicating a likely double bind. The success of double-bind therapy will depend on the properties of the tumour and the treatments used. Finding optimal double-bind strategies (e.g., when to start each treatment; duration; dose) requires calibration of both cell-intrinsic characteristics (e.g., growth rates, costs of resistance) and cell-extrinsic interaction dynamics (e.g., cell competition) within the heterogeneous mix of tumour cell phenotypes being therapeutically targeted. To study these complex dynamics and identify optimal strategies, we developed mathematical models to describe the key cell-intrinsic and cell-extrinsic features of radiation-sensitive, radiation-resistant, and natural killer cells. We calibrated the models using *in vitro* experiments. Cost of resistance was measured in an isogenic prostate cancer cell line model of radiation resistance. Here, cost of resistance is defined as a reduction in growth rate of the resistant line, when compared to its treatment-naïve counterpart. Interaction dynamics (e.g., cooperation or competition between cell types) were quantified using evolutionary game theory (EGT) mathematical models parameterized from *in vitro* competition assays of treatment-naïve isogenic lines in co-culture with RT-resistant lines. Preferential targeting was quantified by extending these competition assays to include NK cells. Taken together, results from our *in vitro* model demonstrate the potential for combining RT and NK cells as an evolutionarily informed treatment strategy in advanced prostate cancer.

## Results

### Radiation resistant cells express higher levels of NK receptor ligands

An isogenic cell line model of RT resistance was previously developed using thirty 2Gy fractionated doses (total 60Gy) on 22Rv1 prostate cancer cells as demonstrated in Figure 2A-B and previously published (24). We measured the surface expression of NK receptor ligands, poliovirus receptor (PVR), poliovirus receptor-related 2 (PVRL2, CD112), and MHC class I polypeptide-related sequence A and B (MICA/B) on control (AMC-22Rv1) and radiation-resistant (RR-22Rv1) cells. PVR and PVRL2 bind activating NK receptors DNAM-1 (CD226) and CD96 or inactivating NK receptor TIGIT (Figure 2G). We found that expression of PVR and PVRL2 was significantly increased on RR-22Rv1 (Figure 2C-D) (p<0.01 and p<0.0001, respectively). NKG2D ligands, MICA and MICB, were upregulated on radiation-resistant cells compared to control (Figure 2C-D) (p<0.01). The mean fluorescence intensity (MFI) for checkpoint ligand Programmed death-ligand 1 (PD-L1) increased significantly in the radiation-resistant cell line (p<0.01) (Figure 2C-D). However, both lines had very low levels (below 400 MFI) of PDL-1 expression. The biological relevance of these low levels remains unknown. Human leukocyte antigen ABC (HLA-ABC, MHC class I) is often upregulated following RT; however, constitutive HLA-ABC was not altered in our isogenic cell line model (Figure 2C-D).

**Figure 2.**
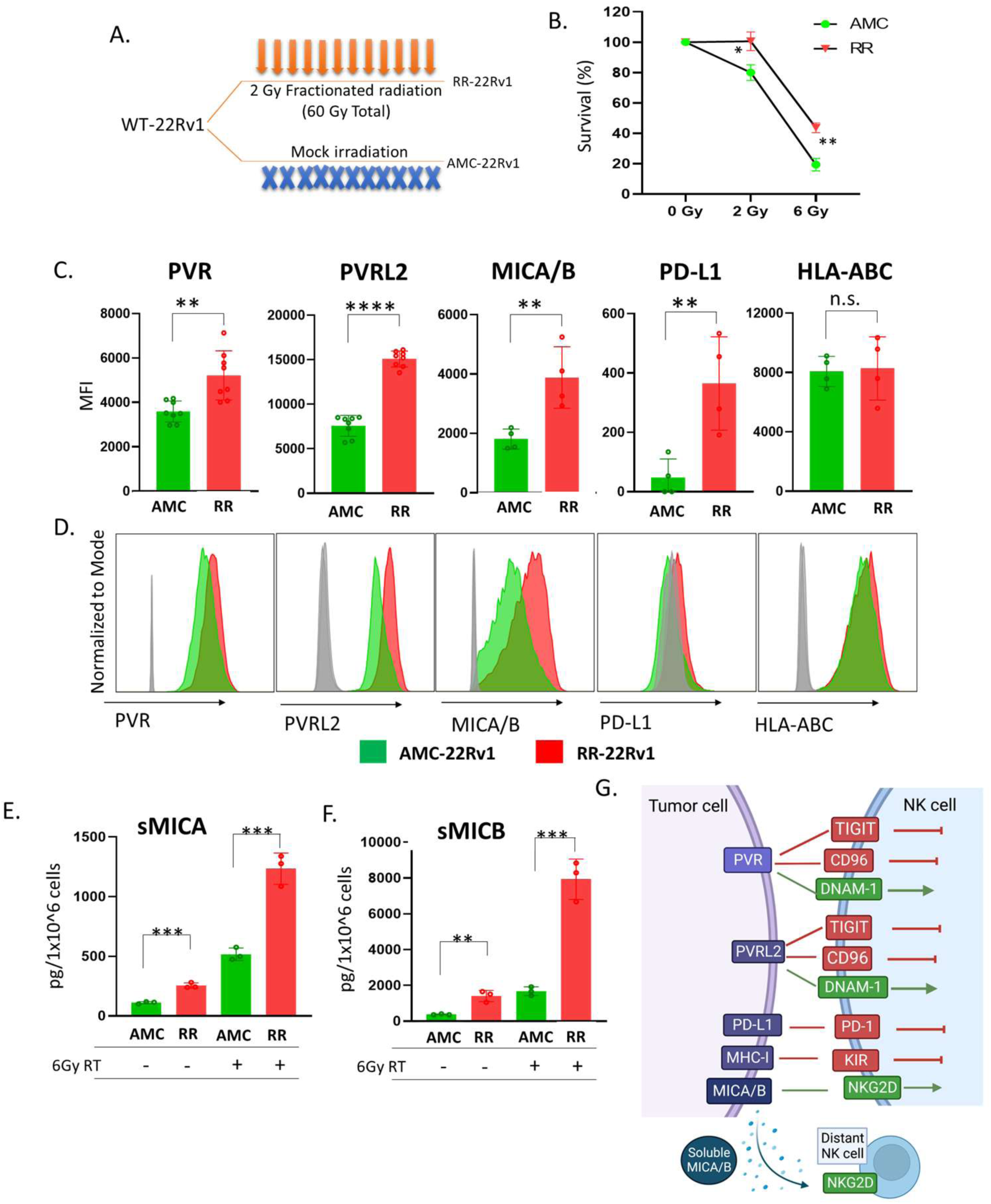
NK ligand expression in an isogenic cell line model of radiation resistance. Prostate cancer cells (WT-22Rv1) were previously exposed to 60Gy (in 2Gy fractions) or mock irradiation to establish an isogenic cell line model with radiation sensitive (AMC) and radiation-resistant (RR) cells lines (**A-B**, (1)). Surface expression of NK ligands by AMC and RR cell lines was measured by flow cytometry, mean fluorescence intensity (MFI) **(C)** and representative histograms (Isotype (grey), AMC (green), RR (red)) **(D)**. Soluble MICA **(E)** and soluble MICB **(F)** were measured by ELISA. Illustration of the effects of NK receptor ligation on NK cell function **(G)**. n.s. = not significant, **p<0.01, ***p<0.001, ****p<0.0001.

Surface expression of MICA and MICB on tumour cells activates NK cells through binding with NKG2D (Figure 2G). Exterior portions of MICA and MICB can be proteolytically cleaved by several metalloproteases resulting in soluble MICA/B proteins (sMICA, sMICB) that maintain full ligand function (Figure 2G). Using enzyme-linked immunosorbent assays (ELISA), we measured the amount of cleaved MICA and MICB in the media of control and radiation-resistant cells before and after radiation. RR-22Rv1 cells have higher levels of sMICA (Figure 2E) and sMICB (Figure 2F) than control lines. Radiation increased levels in both cell lines (Figure 2E-F).

### Radiation-resistant cells are more sensitive to NK cell-mediated killing

NK cell activation occurs when activation signals exceed inhibitory signals from target cells (Figure 2G). Increased NK ligand expression on tumour cells can increase sensitivity to NK cell-mediated killing, however, many NK receptor ligands bind activating and inhibitory receptors on NK cells. Therefore, we directly measured sensitivity to NK cells in our cell lines. AMC-22Rv1 cells and RR-22Rv1 cells were plated in separate wells and treated with RT (6Gy) or co-cultured with an immortalized NK cell line, NKL. Figure 3 illustrates the potential evolutionary relationship between RT-resistance and NK cell killing. Under RT, RR-cells show outpaced growth compared to AMC-22Rv1 cells (Figure 3A). In contrast, increased sensitivity to NK cells by the radiation-resistant line was seen using live imaging (Incucyte, 5:1 E:T ratio) (Figure 3B) and in short term assays using flow cytometry (24 hours, 10:1 E:T) (Figure 3C).

**Figure 3.**
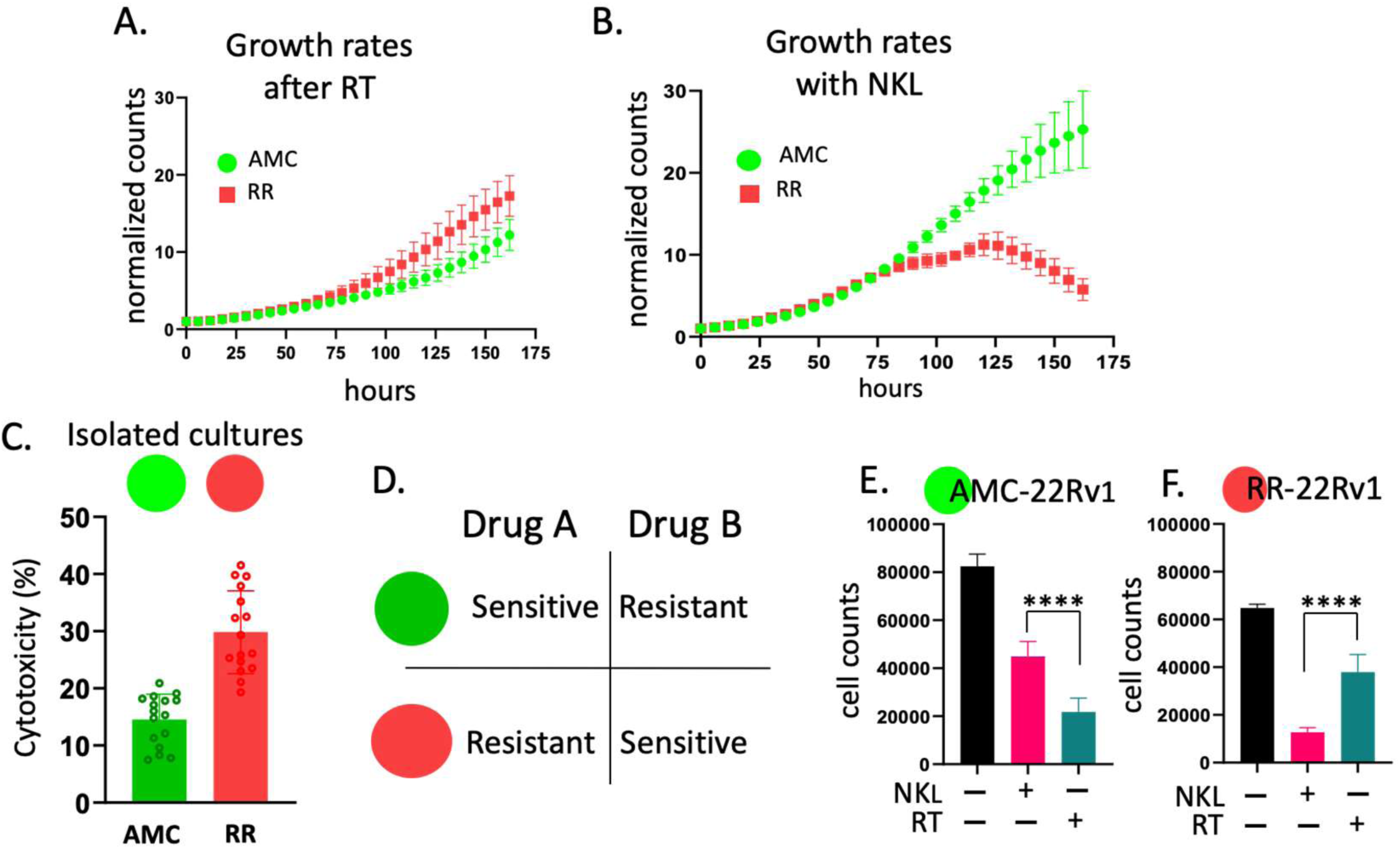
Radiation-resistant prostate cancer cells are more sensitive to NK cell-mediated killing. Live imaging assays of isolated cultures of radiation-sensitive cells (AMC, green) and radiation-resistant cells (RR, red) following radiation (6gy) **(A)** and during challenge with NKL cells (5:1 NK:target cell ratio) **(B)**. Radiation resistant cells (RR) (red) were more sensitive to killing by NKL in a 24-hour flow-based assay (10:1 NK:target cell ratio) **(C)**. Evolutionary double-bind occurs when one treatment targets one population and the second treatment preferentially targets another population **(D)**. Cell counts using Incucyte imager after 7 days of treatment with either 6Gy RT (blue bar) or NKL cells (pink bar) demonstrate that the radiation-sensitive population is inhibited by RT more than NK cells **(E)** and that radiation-resistant cells are inhibited by NK cells more than RT **(F)**. n.s. = not significant, **p<0.01, ***p<0.001, ****p<0.0001

An evolutionary double bind requires differential sensitivity to two treatments. One treatment must strongly inhibit sensitive tumour cells, while a second treatment strongly inhibits cells resistant to the first. Therefore, when the second treatment is applied, cells that were initially sensitive to the first will be selected for, and cells resistant to the first will be selected against (Figure 3D). To determine if NK cells represent an evolutionary double bind with RT in our isogenic cell lines, we compare sensitivity of each line to each treatment (Figure 3E-F). AMC-22Rv1 cells are more sensitive to RT than to NK cells (Figure 3E) while RR-22Rv1 cells are more sensitive to NK cells than to RT (Figure 3F).

### Outcomes are dependent on initial seeding frequency *in vitro*

Treatment sensitivity assays are often performed with populations cultured in separate dishes. This ignores interaction effects occurring between two populations that may affect growth dynamics and treatment sensitivity. Therefore, we repeated the 24-hour flow-based assays in culture dishes seeded with AMC and RR cells in the same well along with NK cells. The NK cells preferentially targeted the RR cells (Figure 4A) similar to when the two tumour cell lines were challenged with NKL cells in isolation (Figure 3C).

**Figure 4.**
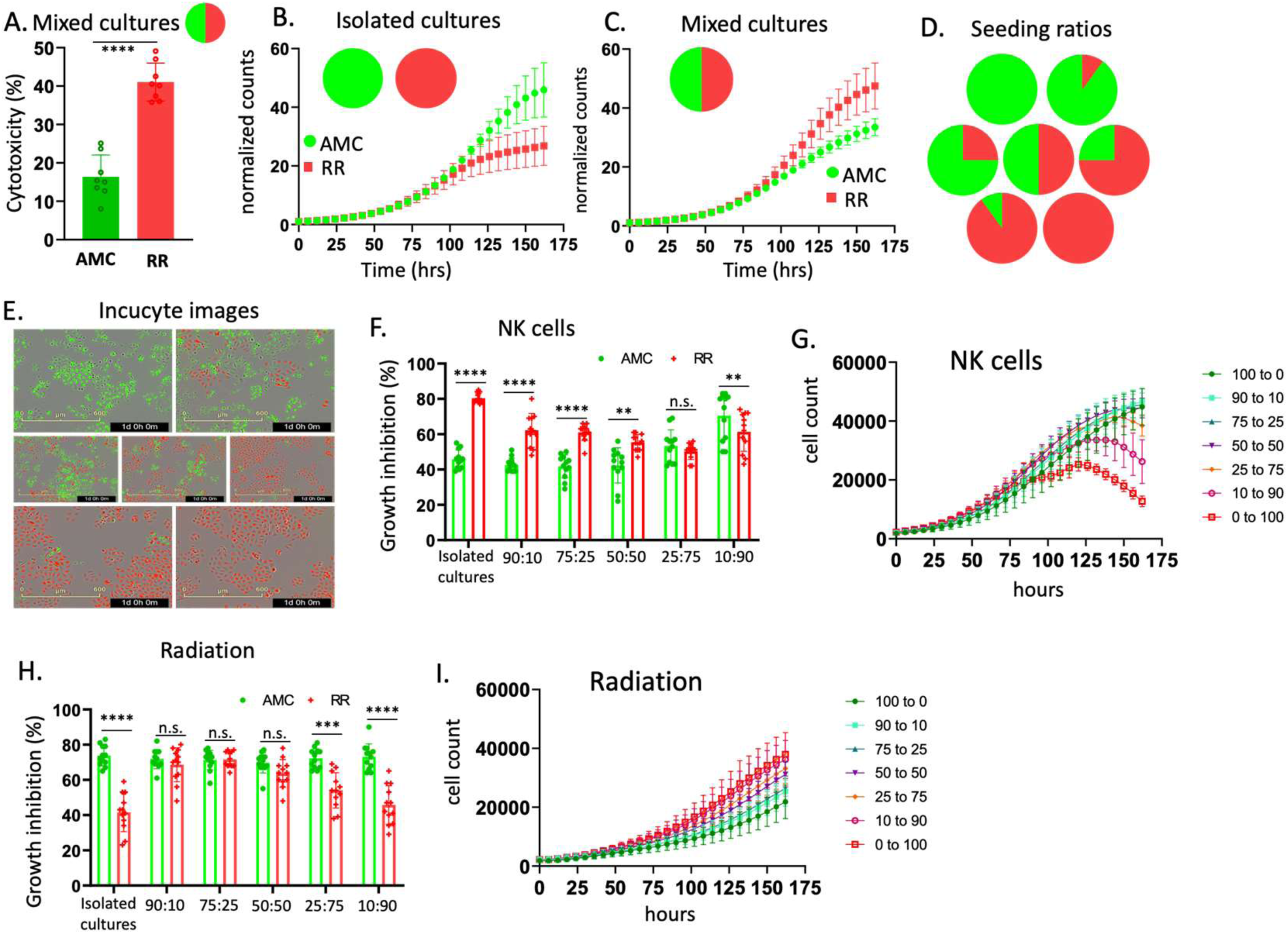
Treatment sensitivity is dependent on initial seeding frequency in mixed cultures. *NKL cells (10:1 effector: target ratio) preferentially target* radiation-resistant cells in mixed cultures (24 hours, 50:50 AMC:RR) **(A)**. AMC and RR cell lines have differential growth dynamics when cultured in isolated wells **(B)** or in mixed cultures (50:50 AMC:RR) **(C)**. Radiation-sensitive (green) and radiation-resistant (red) cells were seeded at various frequencies (0:100, 90:10, 75:25, 50:50, 25:75, 10:90, 0:100 AMC:RR) **(D)** and imaged over 6 days **(E)** in the presence of NKL (5:1 effector: target ratio) **(F-G)** or following radiation (6Gy) **(H-I)**. n.s. = not significant, **p<0.01, ***p<0.001, ****p<0.0001.

Interestingly, the intrinsic growth rates of the two cell lines differ in 50:50 mixed cultures compared to isolated cultures in the absence of treatment (Figure 4B-C), indicating an interaction effect between the two populations where RR-22Rv1 cells inhibit the growth of radiation-sensitive cells and AMC-22Rv1 cells facilitate the growth of radiation-resistant cells. To determine the consequence of these interaction effects on the RT/NK double bind, we cultured fluorescently labelled cells (AMC-22Rv1 GFP, RR-22Rv1 mCherry) at various seeding ratios (Figure 4D-E). We then used live imaging to measure growth rates of the two cell lines under selection with NK cells or single dose RT. NK cells were most effective when RR-22Rv1 cells were cultured alone or seeded at a high frequency (10:90 AMC:RR) (Figure 4F-G, Supplemental Figure 1A). Higher growth inhibition of RR-22Rv1 by RT occurred in cultures with higher numbers of AMC-22Rv1 cells and RT was most effective in cultures with higher frequency of radiation-sensitive cells (Figure 4H-I, Supplemental Figure 1B).

### Two treatments are better than one

Combination therapy is often more effective than monotherapy. Using our *in vitro* isogenic cell line model, we irradiated all seeding ratios (Figure 4D-E) with 6 Gy RT and then added NK cells (5:1 effector to target ratio). NK cells were added after radiation to avoid effects of RT on NKL survival and function. Combination therapy was more effective than either monotherapy in AMC-22Rv1 cells (Figure 5A) and 22Rv1 cells (Figure 5B) in both mono-cultures and mixed cultures (50:50 ratio) (Figure 5C-D). Combination therapy had the largest effect on the radiation-sensitive population. Giving RT prior to NK cells significantly increased the sensitivity of the AMC population to NK cells (Figure 5E) and the addition of NK cells increased the efficacy of RT against the AMC population, although to a lesser degree (Figure 5F). This effect was observed less in the RR-22Rv1 cells (Figure 5E-F). Combination therapy was most effective in all seeding ratios (Supplemental Figure 2).

**Figure 5.**
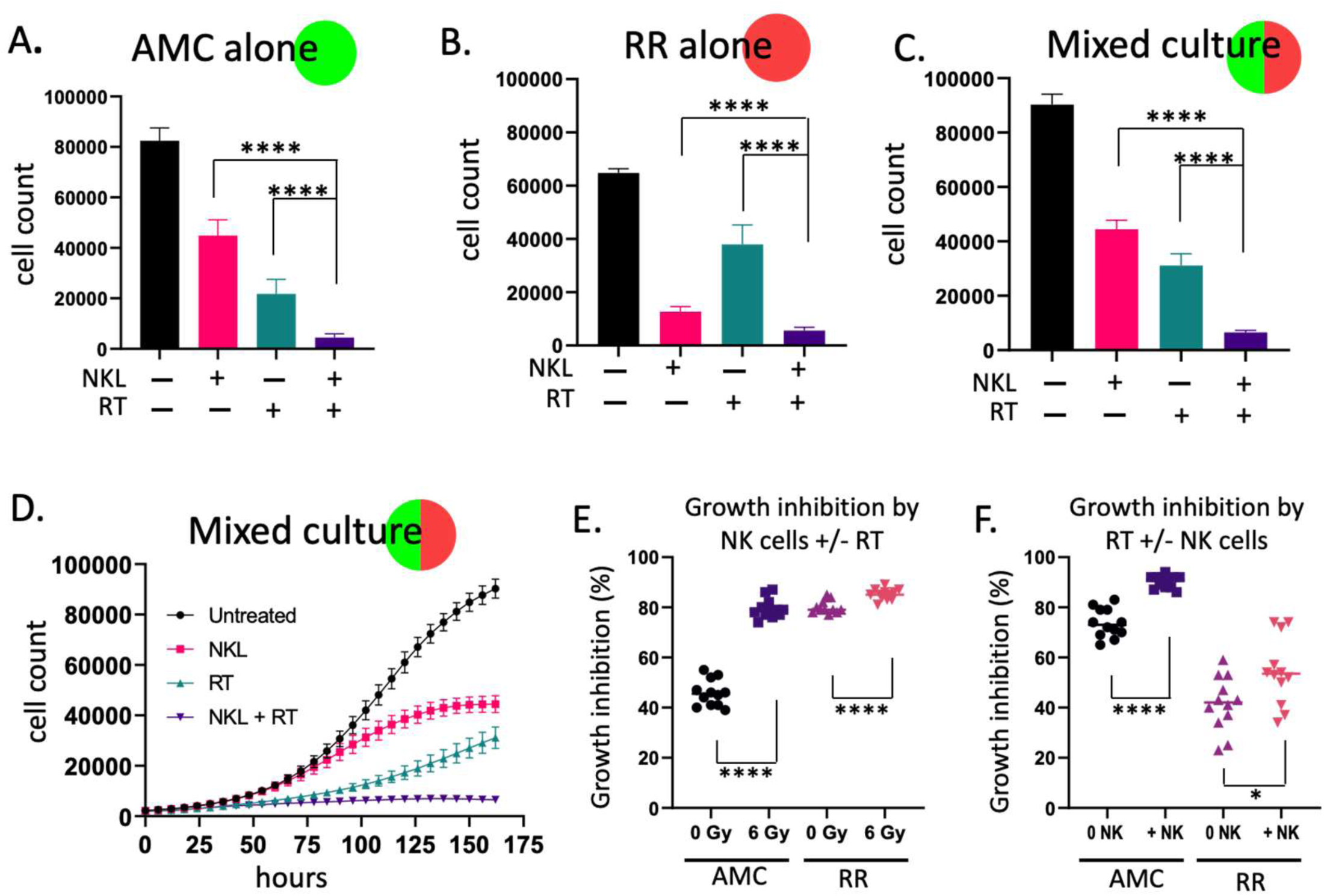
Radiotherapy and NK cells are most effective in combination. Cell counts using an Incucyte imager after 7 days of treatment with NKL cells (5:1 effector to target ratio) (pink bars), 6 Gy RT (blue bars), or combination (purple bars) of radiation-sensitive (AMC) cell line **(A)**, the radiation-resistant (RR) cell line **(B)**, and AMC co-cultured with RR, 50:50 AMC:RR **(C-D)**. Growth inhibition by NK cells with and without RT **(E)**. Growth inhibition by RT with and without NK cells **(F)**.

### Mathematical modelling quantifies an evolutionary double bind

Next, we developed a mathematical model designed to quantify important cell-intrinsic features (e.g., cell-line-specific growth rates) and cell-cell interaction dynamics. The mathematical model (Figure 6E and Supplemental Figure 3) is a Lotka-Volterra (LV) competition model between RT-sensitive (AMC) and RT-resistant (RR). LV and Evolution Game Theory (EGT) mathematical frameworks have been extensively applied to cancer modelling (25) often paired with experimental data to describe competition (26) facilitation (27), or coexistence (28) between cancer cell populations, and to investigate the importance of a cost of resistance (4). Non-cancer cell types (e.g. immune cells) can be considered as a player in the game, or as altering the parameterization of game dynamics between cancer cell types (29, 30).

**Figure 6.**
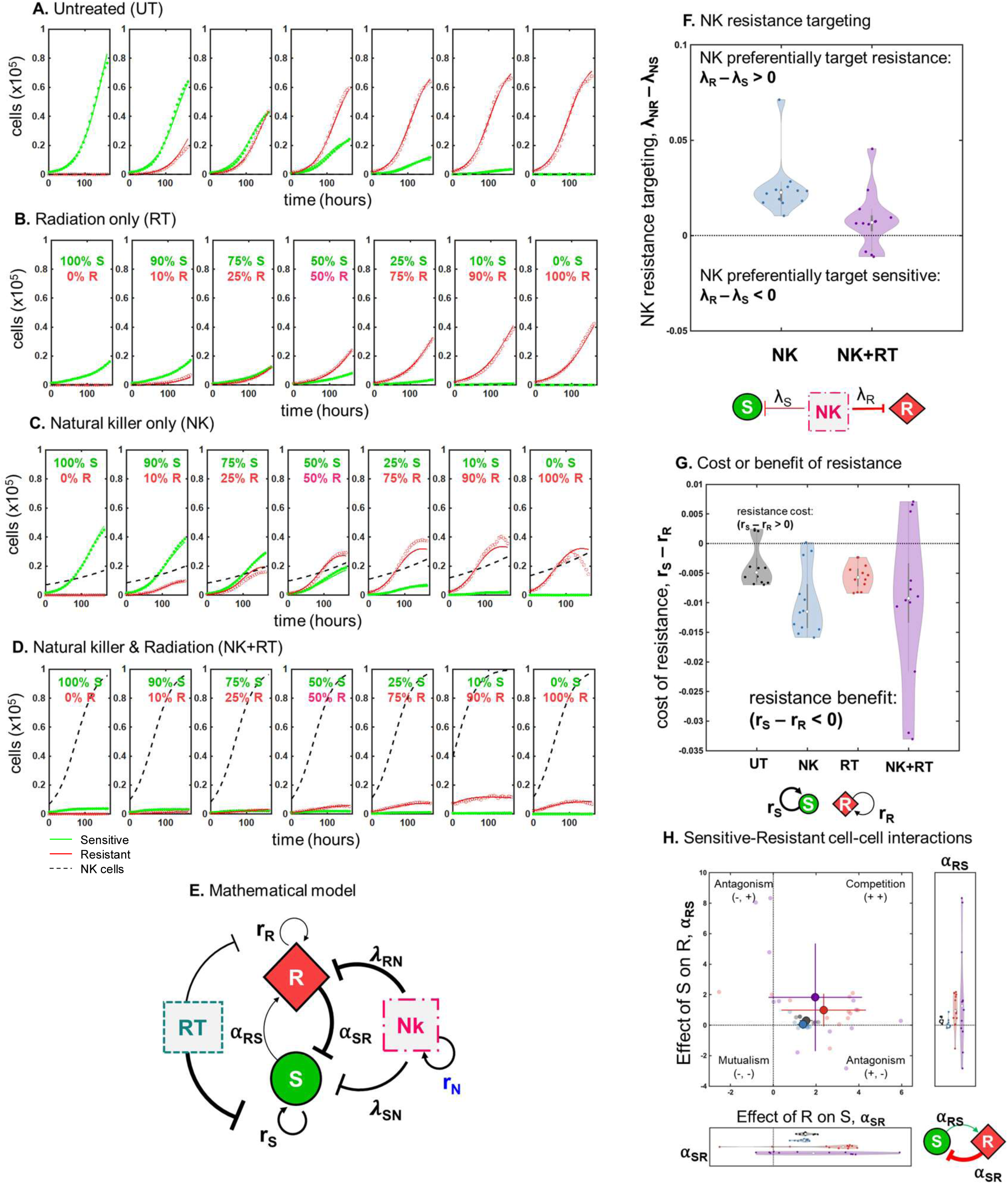
Mathematical model of sensitive and resistant dynamics under therapy. The mathematical model (Eqs. 1-3) was fit across each initial resistant fraction simultaneously and repeated for each of the twelve replicates (see Methods) to give a distribution of parameter values, for untreated **(A)**, RT **(B)**, NK **(C)**, and RT+NK **(D)**. **(E)** Interaction network of the mathematical model. **(F)** When NK cells are present (NK-only or NK+RT), the competitive effect of NK cells on resistant cells (λ_NR_) is stronger than the competitive effect of NK cells on sensitive cells (λ_NS_), indicative of an evolutionary double-bind. **(G)** The cost of radiation resistance (r_S_ – r_R_) is shown to be negative across all treatment conditions, indicating an intrinsic growth benefit in favor of RT-resistant cells. **(H)** Competition dynamics between RT-sensitive and RT-resistant cells indicate “competitive exclusion” where RT-resistant cells outcompete RT-sensitive cells (λ_SR_, λ_RS_ > 0). The effect of R on S is much stronger than that of S on R.

This mathematical model accounts for cell-type-dependent growth rates (r_S_, r_R_), the competitive effect of cell type j on cell type i (⍺_ij_), and cell-type-dependent carrying capacities (K_S_, K_R_). The model also NK cells, which grow with logistic growth dynamics and kill sensitive and resistant cells at a rate of λ_R_ and λ_S_, respectively. The mathematical model was fit to co-culture data (Figure 6A-D) where cells are seeded across a range of initial RT-resistant fractions: f = [0, 0.1, 0.25, 0.50, 0.75, 0.9, 1]. The model was fit to this full range simultaneously and repeated across twelve replicates. Using least-squares error minimization, the model provided good agreement with the data (Figure 6A-D). The distribution of estimated parameters that provided the best fit across replicates is shown in Table 1.

**Table 1:** Parameter estimates under different treatment conditions. Mean values across twelve replicates with plus or minus one standard error measurement shown in parentheses. See equations 1 – 3.

| | $r_S$ | $r_R$ | $r_N$ | $K_S$ | $K_R$ | $K_N$ | $\alpha_{SR}$ | $\alpha_{RS}$ | $\lambda_{SN}$ | $\lambda_{RN}$ |
| --- | --- | --- | --- | --- | --- | --- | --- | --- | --- | --- |
|  | growth rate (S) | growth rate (R) | growth rate (N) | carrying capacity (S) | carrying capacity (R) | carrying capacity (N) | competition effect of R on S | competition effect of S on R | NK kill rate of S cells | NK kill rate of R cells |
| UT | **0.032**<br>(0.0315, 0.0325) | **0.0357**<br>(0.035, 0.0363) | **0**<br>(fixed) | **1.27**<br>(1.24, 1.31) | **0.779**<br>(0.757, 0.8) | **1**<br>(fixed) | **1.54**<br>(1.45, 1.63) | **0.3**<br>(0.254, 0.347) | **0**<br>(fixed) | **0**<br>(fixed) |
| NK | **0.034**<br>(0.033, 0.0349) | **0.0438**<br>(0.0425, 0.045) | **0.0178**<br>(0.0162, 0.0193) | **1.27**<br>(fixed) | **0.779**<br>(fixed) | **1**<br>(fixed) | **1.38**<br>(1.3, 1.46) | **0.0526**<br>(-0.0334, 0.139) | **0.0308**<br>(0.0242, 0.0374) | **0.0558**<br>(0.0471, 0.0645) |
| RT | **0.0162**<br>(0.0158, 0.0167) | **0.0218**<br>(0.0211, 0.0225) | **0**<br>(fixed) | **1.27**<br>(fixed) | **0.779**<br>(fixed) | **1**<br>(fixed) | **2.36**<br>(1.79, 2.93) | **0.99**<br>(0.68, 1.3) | **0**<br>(fixed) | **0**<br>(fixed) |
| RT+NK | **0.023**<br>(0.0199, 0.0262) | **0.0331**<br>(0.0268, 0.0394) | 0.0245<br>(0.0204, 0.0286) | **1.27**<br>(fixed) | **0.779**<br>(fixed) | **1**<br>(fixed) | **1.96**<br>(1.33, 2.59) | **1.83**<br>(0.809, 2.85) | **0.0865**<br>(0.0488, 0.124) | **0.0945**<br>(0.0545, 0.135) |

The mathematical model allowed us to evaluate our hypothesized existence of a double-bind by comparing parameters describing the inhibition of radiation-sensitive (λ_SN_) and radiation-resistant (λ_RN_) cells by NK cells. NK cells preferentially target RT-resistant cells (and thus represent a true double bind) if the difference in these inhibition rates is positive i.e. λ_RN_ – λ_SN_ > 0. Figure 6F confirms the existence of the double bind in NK cell therapy alone (blue) and in combination with RT (purple): λ_RN_ – λ_SN_ > 0. This effect is most pronounced with NK therapy alone but is still present with combination therapy.

### The double bind does not require a cost of resistance

Next, we investigated whether a cost to radiation-resistance exists in terms of growth rate differences between sensitive and resistant populations (r_S_ > r_R_). In Figure 6G, model analysis indicates a lack of cost of resistance—indeed, an intrinsic growth benefit is conferred to RT-resistant cells, across all treatment conditions (r_R_ > r_S_). However, this benefit is offset by a lower carrying capacity (see Table 1) such that K_R_ < K_S_. Interestingly, it appears that a cost of resistance to intrinsic growth rate is not a necessary condition for an evolutionary double bind.

Finally, we quantified the competitive effects of RT-sensitive cells on RT-resistant cells (⍺_RS_) and the inverse (⍺_SR_). RT-sensitive cells have a weak effect on RT-resistant cells (⍺_RS_ ≈ 0) while RT-resistant cells strongly suppress RT-sensitive cells (⍺_SR_ > 0). These competition parameters are plotted in Figure 6H to classify interactions between sensitive and resistant populations into four possible categories: mutualism, competition, sensitive antagonism, or resistant antagonism.

This confirms previous results of competitive dynamics in alternative radiation-sensitive and resistant cell lines (31). Again, it appears that a frequency-dependent fitness disadvantage of RT-resistant cells is not a necessary condition for an evolutionary double bind.

### Clinical relevance of an evolutionary double bind

Next, to illustrate the clinical relevance of a second drug that fits the definitions of an evolutionary double-bind, we developed a simplified mathematical model with the following components: logistic growth, shared carrying capacity, a cost of resistance, and a double-bind parameter, as seen in equations 4 – 5 (materials and methods). The double-bind parameter (B) modifies the second drug’s target by either only targeting sensitive cells (B=0), only targeting resistant cells (B=1), or as a ratio of efficacy against sensitive and resistance cells (0 < B < 1).

The simplified model is used to gain intuition about the role of preferential targeting by the secondary drug in an evolutionary double-bind. Figure 7 shows that an increasing dose of radiation therapy reduces the sensitive population (figure 7B, top) and selects for resistance (figure 7B, bottom). Figure panels 7C – E illustrate the importance of the double-bind parameter when considering a second drug as a candidate for combination therapy. When the evolutionary double-bind is weak (B ≈ 0), the effect of adding this second drug only serves to further reduce the sensitive population (figure 7C), while not forestalling the emergence of resistance. In contrast, when the double-bind is strong (B ≈ 1), the second drug preferentially targets the resistant population. In general, the final sensitive population size increases with B, while the final resistant population size decreases with B (figure 7E). Thus, a viable clinical strategy is to screen for drugs with a strong evolutionary double-bind to suppress resistance to front-line therapies, which in turn strongly target treatment-sensitive populations.

**Figure 7.**
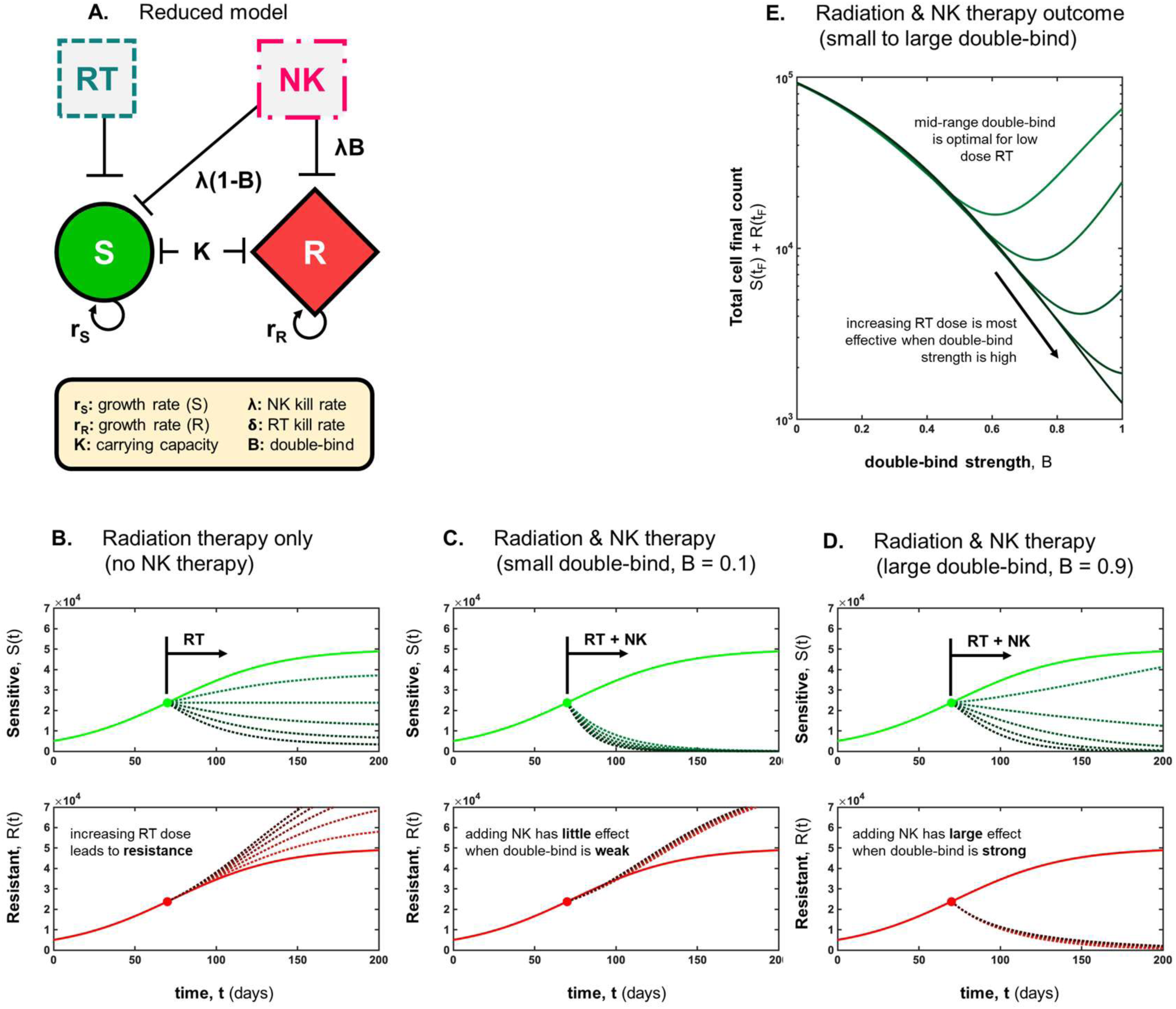
Reduced mathematical model. (**A**) The mathematical model is reduced to its baseline components: logistic growth, shared carrying capacity, and a double-bind parameter, as seen in equations 4 – 6. **(B)** Increasing radiation therapy reduces the sensitive population (top) while selecting for resistance (bottom) Increasing values of δ shown in darker colors; δ = [0.5, 1, 1.5, 2, 2.5]). **(C)** When double-bind is weak (B = 0.1) the addition of NK therapy further reduces sensitive population, but has little effect on resistance. **(D)** When double-bind is strong, addition of NK therapy reduces resistant population, and sensitive population can be controlled through radiation. (E) Full range of double-bind parameter, 0 ≤ *B* ≤ 1. Parameters unless otherwise noted: λ = 0.05; K = 1e5.

## Discussion

Evolution of resistance remains a substantial barrier to control and cure of clinical metastatic cancers by single treatments applied using the traditional oncologic strategy of continuous treatment at maximum tolerated dose (MTD) until progression (9, 32, 33, 34). Typically, second-line therapy is less effective than front-line treatment, in part because of cross-resistance to first-line therapy results in some protection to subsequent lines of therapy. In contrast, double-bind therapy anticipates the cancer cell’s adaptive response to an initial therapy and then specifically exploits that strategy with the follow-on treatment (7). Thus, the double-bind strategy is designed to *increase* the efficacy of second-line therapy. Ideally, these dynamics result in second-line treatment that is more effective than initial treatment. Importantly, therapies that target resistant strategies to front-line drugs are often less effective when given first. As a result, identifying optimal second-line therapies requires a detailed understanding of the underlying evolutionary dynamics, since their efficacy as isolated monotherapies may obscure their potential as a double-bind therapy.

Here, we utilized a combined empirical and theoretical investigative strategy to understand the eco-evolutionary dynamics affecting double-bind therapy optimization. Specifically, we demonstrate that radiation resistance alters NK cell ligand expression and increases sensitivity to NK cell-mediated killing in an isogenic cell line model. Radiation induces activation of double-stranded DNA break repair pathways, homologous recombination (HR), and non-homologous DNA end joining (NHEJ) (24, 35, 36, 37, 38). Increased expression of repair pathway proteins (DNA-PK, ATM, Rad52, MLH1, and BRCA1) is associated with cancer cell survival following radiation (24, 39). Therefore, NK cell-based immunotherapies are an obvious choice for combination with DNA damaging agents for a double-bind strategy.

We have demonstrated an effective double-bind therapy using RT and NK cells in an *in vitro* cell line. These findings will need to be tested in additional model systems before clinical relevance can be determined. Pre-clinical methods for identifying potential double-bind therapies have not been fully elucidated. Like *in vitro* models used to identify synergistic effects of combination therapies, ‘gold-standard’ experimental procedures will need to be defined for high-throughput identification of double-bind combination therapies. Our findings provide an experimental setup that highlights the need for mixed cultures in double-bind studies.

To our knowledge, this manuscript provides the first direct experimental evidence quantifying a treatment double bind in prostate cancer. Our work is also mathematically novel in that it extends Evolutionary Game Theory models to consider the competitive effect of NK cells on treatment-naïve and RT-resistant lines, thus adding a new dynamic to the double-bind concept: preferential targeting of RT-resistance. These mathematical models can also be used with other *in vitro* model systems and can be extended to include additional therapy considerations such as dose, timing, and additional treatment modalities.

The theoretical literature often assumes *a priori* a fitness cost of adaptive resistance mechanisms (40, 41). Here, we investigated both an intrinsic fitness cost to growth rates and a frequency-dependent fitness advantage and found neither. In fact, we observed a growth rate benefit to resistant cells, albeit at a cost to their carrying capacity, and suppression (via competitive exclusion) of sensitive cells. Previously proposed evolution-based treatment strategies, such as adaptive therapy, have shown that a cost of resistance is necessary pre-requisite (41, 42, 43). In contrast, the evolutionary double-bind treatment strategy of NK cell-based therapy represents a promising alternative that neither requires a cost of resistance or dose modulation but rather focuses on the critical order of treatment.

## Materials and methods

### Cell lines

Isogenic cell lines AMC-22Rv1 and RR-22Rv1 were obtained from Laure Marignol (Trinity St. James’s Cancer Institute) and were previously derived from parental line 22Rv1 with either fractionated radiation (30 x 2Gy) or mock irradiation (24). The resulting cell lines, radiation-resistant (RR-22Rv1) and aged matched control (AMC-22Rv1), have differential sensitivity to X-ray radiation. Cells were maintained in RPMI (Gibco) with 10% fetal bovine serum (FBS) and 1% pen/step in a humidified incubator with 5% CO2 at 37DC. Cells were kept in log phase except were specified. NKL cells were obtained from internal lab stocks and cultured in RPMI with 10% FBS, 1% pen/strep, and 100 IU recombinant human IL-2. Cell lines were authenticated and mycoplasma tested regularly (Lonza).

Stable GFP and mCherry expressing cell lines were created through lentiviral transduction with pLV(Exp)-CAG>3xNLS/EGFP-P2-A-puro and pLV(Exp)-CAG>3xNLS/mCherry-P2-A-puro, respectively. Lentivirus-EGFP and –mCherry were kindly provided by Marusyk Lab (Moffitt Cancer Center). Positively transduced cells were selected with 2 µg/ml puromycin followed by a sterile sort for high GFP and mCherry expression (BD FACSAria II). Expression was maintained with 1 µg/ml puromycin in all experiments except where immune cells were needed.

### Flow Cytometry

Following treatment (where applicable) tumour cell lines were stained with Zombie Aqua Fixable viability dye (1:1000 in PBS) and for NK ligand (Biolegend) expression in PBS containing 5% FBS and 0.1% sodium azide for 20-30 min at room temperature. Secondary antibody was added for ULBP 1 and 2/5/6 only (R&D Systems). Cells were then washed and fixed in 4% paraformaldehyde. Flow was performed on BD LSR II (Trinity School of Biochemistry and Immunology or Moffitt Cancer Center) and analysed using FlowJo software (Tree Star). Internal controls were used in all experiments and technical and biological replicates were always performed in the same facility to remove inter-machine variability. Morphology gating, viability gating, and doublet discriminator gates were used prior to calculating mean fluorescence intensity (MFI). Mean fluorescence intensity was determined as MFI stained – MFI unstained. ULBP primary and secondary antibodies were purchased from R&D Systems, all others all others were pre-conjugated and obtained from BioLegend.

### Radiation

*In vitro* radiation was given in either 2, 6, 10, or 15 Gy doses using XRAD 160 biological irradiator (Precision X-Ray Inc). Prior to RT cells were plated in complete media with 10% FBS and 5% Pen/Strep and left to adhere overnight. Following RT cells were immediately returned to incubator with 5% CO2 at 37°C.

### Competition experiments with Incucyte imager

Cells were counted and mixed at various ratios prior to plating in 96-well flat bottom plates (Corning). Tumour cells were plated at 2,000 cells per well and left to adhere. When NKL and radiotherapy were given together RT was performed first to avoid RT effects on NKL cell function. Images were taken every 6 hours for 150 hours on the Incucyte ZOOM. Cell number was determined by the Incucyte software. Three technical replicates were used for each run. Normalized cell counts were calculated to allow for growth rate comparison between the different ratios (time t/time 0).

### Statistical analysis

Prism (version 9, GraphPad software La Jolla California) was used for all statistical analysis. A Student t-Test, One-way ANOVA with Bonferroni correction, and two-way ANOVA was used when the means of more than two groups were compared. All experiments were performed with three technical replicates. A minimum of 10,000 events were collected for each flow sample. n.s. = not significant **p<0.01, ***p<0.001, ****p<0.0001.

### Mathematical model development

Below, we introduce a two-population Lotka-Volterra competition model, consisting of sensitive (S), radiation-resistant (R). Both cell types compete for space and resources according to a fixed carrying capacity, K_i_, and a cell-type specific growth rate, r_i_, where i=[S,R,N]. The inter-specific competition terms are given by ⍺_ij_, describing competition of cell type *j* on cell type *i*.

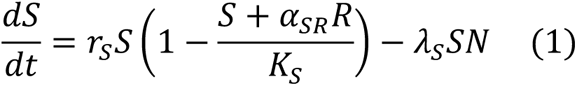

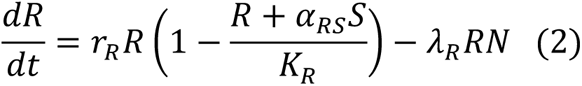

In our experimental system, we make the simplifying assumption that NK cells grow with a logistic growth dynamic, with no competitive feedback from S or R cells:

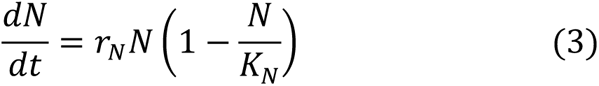

### Fitting process

The following procedure was used when fitting longitudinal experimental data. The dataset contains cell counts for sensitive, S(t), and radiation resistant, R(t), cells over time every six hours over approximately 150 hours, repeated for 12 replicates. Imaging and fluorescence do not give any information about N(t), so we set K_N_=1×10^5^. Let f be the initial fraction of radiation resistant cells in the experimental setup such that a “fitting batch” represents a single replicate for f=[0, 0.10, 0.25, 0.50, 0.75, 0.90, 1]. Data for this single replicate are fit to the model simultaneously across all values of f and subsequently repeated for each of the twelve replicates. In this way, each parameter (r_S_, r_R_, …) has a mean value and standard deviation.

Fitting is done for 1) untreated, 2) radiation treatment only, 3) NK cells only, and 4) combination. Due to concerns about the identifiability of the carrying capacity in the treated experimental conditions, the untreated condition is fit first, and all other treated experimental conditions are fit using the mean value for carrying capacities derived in the untreated condition (K_S_ and K_R_). Growth rates (r_i_) are bound between 0 and 1, competition values (λ_ij_) are bound between –15 and 15, and carrying capacities (K_i_) are bound between 0 and 10^6^. For experimental setups 1 and 2 with no NK cells, the effect of NK cells on S and R is set to zero: λ_S_= λ_R_=0.

### Reduced model

In figure 7, the mathematical model is reduced to its baseline components: intrinsic growth rates, r_S_ and r_R_, shared carrying capacity, K, RT kill rate, 6, NK cell kill rate, λ, and a double-bind parameter, B. Radiation therapy reduces the growth rate of sensitive cells only, by a fraction δ.

NK cells kill cancer cells at a rate λ, where the double-bind parameter alters the second drug’s kill effect to target only sensitive cells (B = 0), target only resistant cells (B=1), or a given ratio of sensitive and resistance cells (0 < B < 1). A cost of resistance reduces the growth rate of resistant cells by a fraction c.

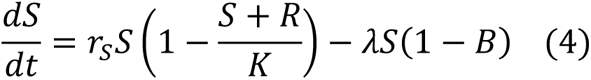

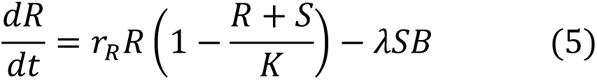

where radiation therapy reduces the growth rate of sensitive cells during therapy by a fraction of (1 – δ):

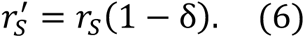

## Supplemental Material

**Supplemental Figure 1.**
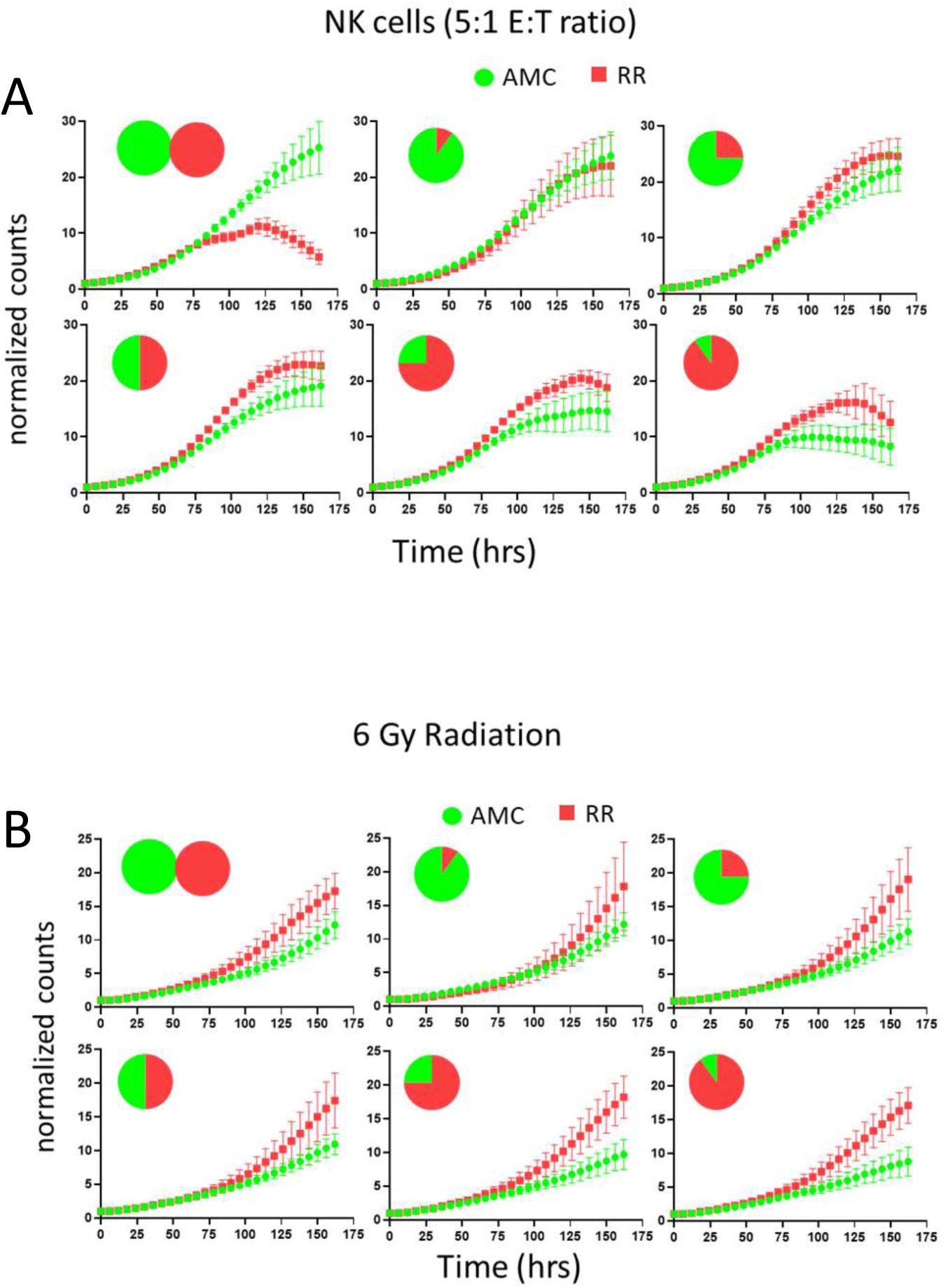
Normalized growth rates of AMC (green) and RR (red) cells treated with NKL cells or 6 Gy RT. Counts are normalized to time 0.

**Supplemental Figure 2.**
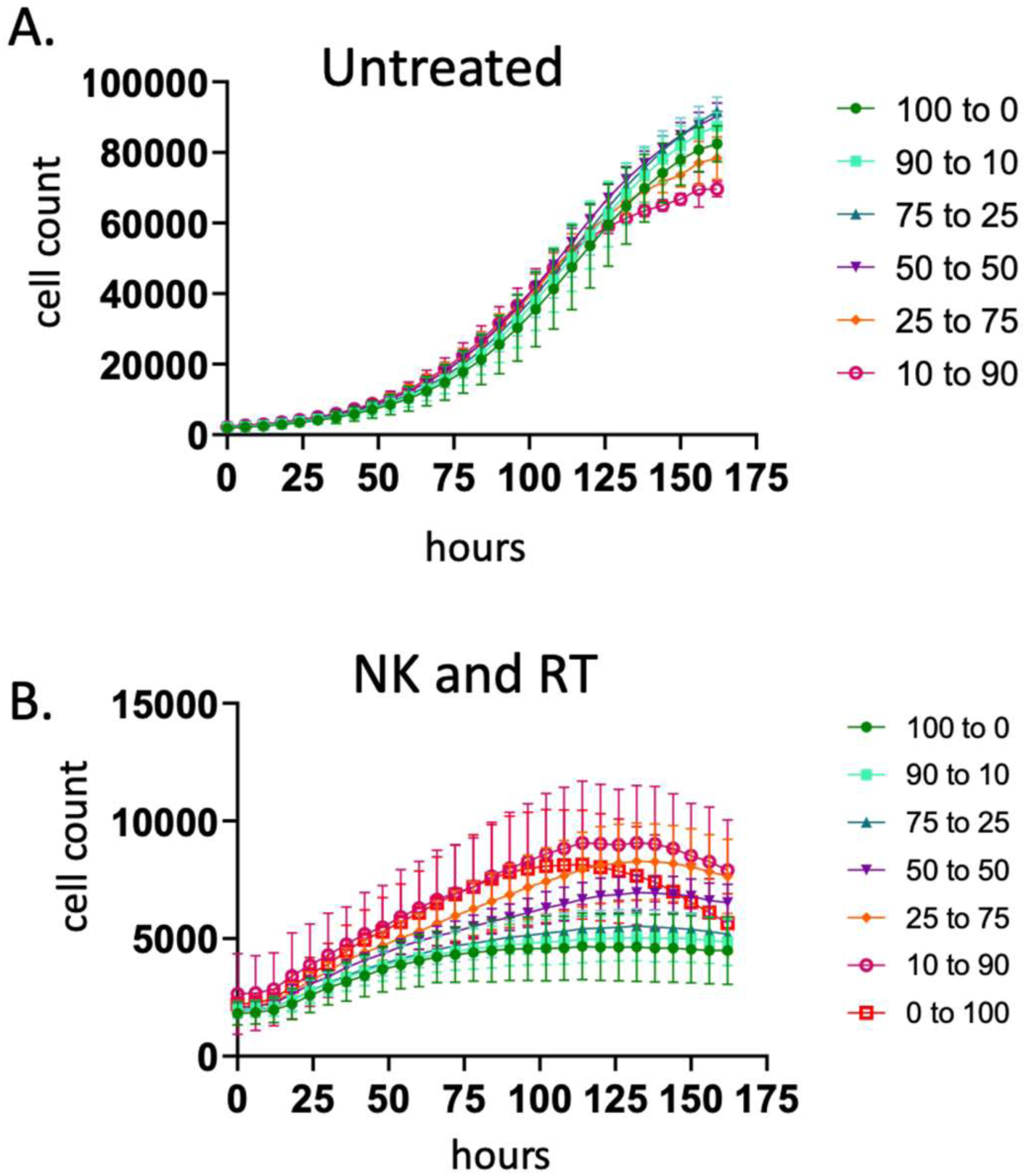
Incucyte data of all seeding ratios untreated (A) or treated with combination therapy, 6 Gy RT followed by 5:1 NK:target cell ration (B).

**Supplemental Figure 3.**
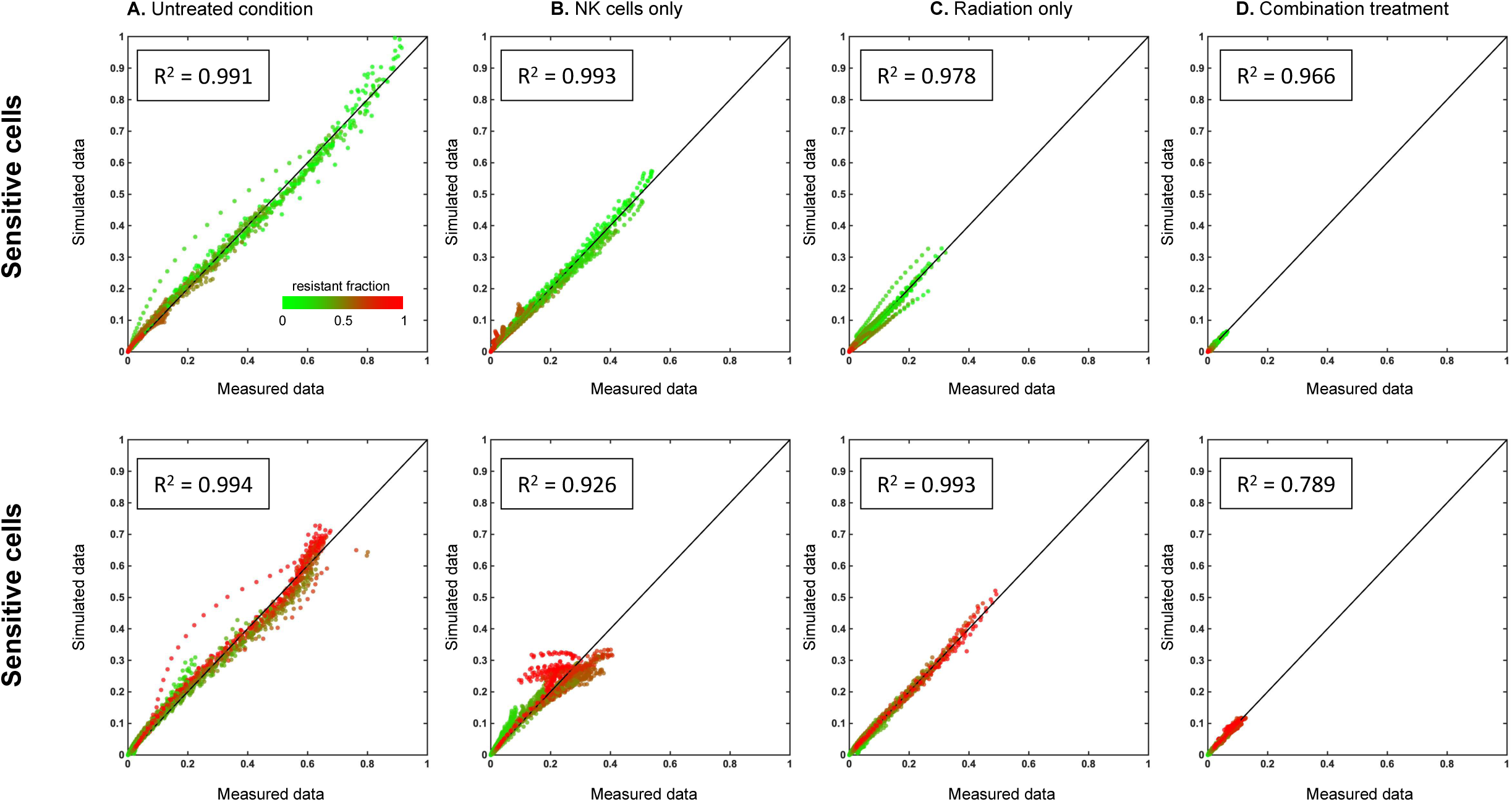
Agreement between the measured data (x-axis) and the mathematical model simulated data (y-axis) for (A) untreated conditions, (B) natural killer cells only, (C) radiation therapy only, and natural killer cells with radiation therapy, repeated for sensitive (top row) and resistant (bottom row) cells.

